# Sex-dependant differences in the ability of nicotine to modulate discrimination learning and cognitive flexibility in mice

**DOI:** 10.1101/2024.04.29.591794

**Authors:** Yoshiatsu Aomine, Yuto Shimo, Koki Sakurai, Mayuka Abe, Tom Macpherson, Takaaki Ozawa, Takatoshi Hikida

## Abstract

Nicotine, an addictive compound found in tobacco, functions as an agonist of nicotinic acetylcholine receptors (nAChRs) in the brain. Interestingly, nicotine has been reported to act as a cognitive enhancer in both human subjects and experimental animals. However, its effects in animal studies have not always been consistent, and sex differences have been identified in the effects of nicotine on several behaviors. Specifically, the role that sex plays in modulating the effects of nicotine on discrimination learning and cognitive flexibility in rodents is still unclear. Here, we evaluated sex-dependent differences in the effect of daily nicotine administration at various doses (0.125, 0.25, and 0.5 mg/kg) on visual discrimination (VD) learning and reversal (VDR) learning in mice. In male mice, nicotine significantly improved performance in VDR, but not VD, task, while, in female mice, nicotine significantly worsened performance in the VD, but not VDR, task. Next, to investigate the cellular mechanisms that underlie the sex differences in the effects of nicotine on cognition, transcriptomic analyses were performed on prefrontal cortex tissue samples from male and female mice that had undergone VD and VDR tasks. Pathway enrichment analysis and Protein-protein interaction (P-PI) analysis using gene sets with altered gene expression found three types of effects of nicotine: those common to both sexes, those in males only, and those in females only. Decreased expression of postsynaptic-related genes in males and increased expression of innate immunity-related genes in females were identified as possible molecular mechanisms related to sex differences in the effects of nicotine on cognition in discrimination learning and cognitive flexibility.

## 2 Introduction

Nicotine, a highly addictive alkaloid contained in tobacco (Gould, 2023; JARVIK, 1991; Mcgrath-Morrow et al., 2020; Zeidler et al., 2007), is known to exert various physiological effects via its agonism of nicotinic acetylcholine receptors (nAChRs), heterogeneous cationic channels that are widely expressed in both neural and non-neuronal tissues (Gotti et al., 2006; Hollenhorst & Krasteva-Christ, 2021; Zoli et al., 2018). In the brain, nicotine binding at nAChRs expressed at presynaptic and preterminal sites promotes the release of neurotransmitters, including dopamine, which regulate widespread brain functions including cognition, reward, learning, memory, anxiety, and appetite (Benowitz, 2008; Kim & Picciotto, 2023).

Past studies in humans and rodents have demonstrated that nicotine can improve cognitive function. In humans, a meta-analysis reported positive effects of nicotine or smoking on attention, as well as short-term episodic and working memory (Heishman et al., 2010), while several other studies have similarly demonstrated enhanced performance on tests of continuous attention, computational processing, and memory (DeVito et al., 2014; Myers et al., 2008; Warburton et al., 1992). In rodent studies, nicotine improved VD in rats and three mouse strains (F. Bovet-Nitti, 1966, 1969), and facilitated reversal learning in a probabilistic reversal learning tasks (Milienne-Petiot et al., 2018). However, conversely, nicotine has also been reported to impair reversal learning in VD tasks, suggesting that nicotine may not improve cognitive function indiscriminately (Cole et al., 2015; Ortega et al., 2013).

Interestingly, there are sex differences frequently observed in the effects of nicotine. In humans, females are typically quicker to develop nicotine dependence, are more likely to demonstrate a depressive tendency during dependence, and have a stronger negative affective response during nicotine withdrawal compared to males (Hogle & Curtin, 2006; Komiyama et al., 2018; Pogun et al., 2017). In experimental animals, female rodents exhibit greater nicotine self-administration behavior (Chaudhri et al., 2005) and behavioral sensitization (Booze et al., 1999; Harrod et al., 2004), and demonstrate a stronger anxiolytic response to acute nicotine (Cheeta et al., 2001) or during nicotine withdrawal (Caldarone et al., 2008). Despite such knowledge, little is known about potential sex differences of the effects of nicotine on cognitive abilities, including VD learning and cognitive flexibility. Thus, examination of such differences using animal studies is necessary for elucidation of the modulatory role of nicotine on cognition.

Among nAChR subunits, the α7 subunit is known to be necessary for nicotinic enhancement of discrimination learning, and positive allosteric modulation of α7 subunit containing nAChRs enhance recognition memory and cognitive flexibility (Milienne-Petiot et al., 2018; Nikiforuk et al., 2015). Alternatively, the β2 subunit has been implicated in the ability of nicotine to impair cognitive flexibility (Cole et al., 2015). Interestingly, transcript levels and the number of binding sites on nAChR subunits in the brain are sexually dimorphic (Bagdas et al., 2019; Cross et al., 2017; Moen & Lee, 2021). Furthermore, nicotine acting at nAChRs is known to result in sex-dependent changes in gene expression in brain regions that have been linked with cognitive control, including the frontal cortices (Friedman & Robbins, 2022). Indeed, chronic nicotine administration was found to result in an up-regulation of the sphingolipid metabolism-related gene *CERKL* in the frontal lobe of male rodents, but a down-regulation in female rodents (Vargas-Medrano et al., 2023). Similarly, gestational nicotine exposure was demonstrated to increase and decrease expression of major myelin genes in the prefrontal cortex (PFC) of male and female mice, respectively (Cao et al., 2013). Although such studies have identified alterations in the expression levels of individual genes, large-scale unbiased investigation of sex-dependent differences in nicotine-induced alteration of gene expression has not been conducted so far.

Here, we investigate the effects of nicotine on discrimination learning and cognitive flexibility in male and female mice. VD learning and VD reversal (VDR) learning tasks were chosen as behavioral tests because of the high translational potential of this experimental platform between rodent and human studies. Furthermore, Bussey-Saksida Touch Systems were used due to their ability to measure complex cognitive functions with minimal researcher interference (Horner et al., 2013; Macpherson & Hikida, 2018; Nishioka et al., 2023). Additionally, we investigate gene expression changes in the PFC induced by short-term or long-term administration of nicotine in male and female mice, in order to help elucidate the molecular mechanisms that may contribute to the ability of nicotine to modulate cognitive abilities and its difference between the sexes.

## 3 Materials and methods

### 3.1 Animals

Male (n=12) and female (n=12) C57BL6/JJcl mice obtained from CLEA Japan Inc (Tokyo, Japan) and aged between 8-10 weeks were used for all experiments. Mice were housed on a 12-hour light/dark cycle (Light: 0800-2000, Dark: 2000-0800) in a quiet environment with room temperature maintained at 24 °C ± 2 °C and 50 ± 5% humidity. Mice were housed according to sex and in groups of between 2-5 mice with *ad libitum* access to food and water until behavioral experiments. All animal experiments complied with institutional guidelines set by Osaka University Institute for Protein Research Animal Research Committee.

### 3.2 Drugs

Nicotine solution was prepared by dissolving nicotine hydrogen tartrate salt (SML1236, Sigma-Aldrich, Missouri, USA) in saline (Otsuka normal saline, Otsuka Pharmaceutical Factory, Inc., Tokushima, Japan). 0.5 mg/kg solution was prepared by dissolving 0.5 mg of nicotine hydrogen tartrate salt in 10 mL of saline; 0.25 mg/kg and 0.125 mg/kg solutions were prepared by diluting the 0.5 mg/kg solution two- and four-fold with saline. The solutions were administered to mice at a volume of 0.1 mL per gram of mouse weight. Doses of nicotine were selected based on those previously reported to facilitate VD in mice and rats (F. Bovet-Nitti, 1966, 1969).

### 3.3 Touchscreen operant chambers

Pre-training and testing were conducted in touchscreen operant chambers (Model 80614, Campden Instruments Ltd., Loughborough, UK). A partition plate with two square holes (W:7 cm, H:7.5 cm) in the center separated by a 5 mm space in between was fitted in front of an infrared touch screen to create two distinct panels for touch responses. A reward tray connected to polyvinyl tube and a peristaltic pump for liquid reward delivery was placed on the opposite side of the chamber. The chamber was always kept dark during experiments, with only three light sources: a touch screen light (panel lights), a light on the top of the reward presentation dish (dish light) and house light. The touchscreen operant chambers were controlled using ABET II (Lafayette Instrument Co., Indiana, USA) and Whisker (Cambridge University Technical Services Ltd., Cambridge, UK) software. Between each task sessions, the chamber was cleaned with 70 % ethanol.

### 3.4 Pre-training

Visual discrimination (VD) Tasks in the Touchscreen Operant System were conducted as previously described with minor modification (Horner et al., 2013; Morita et al., 2016), It consisted of six phases: pre-training (*Initial touch training, Must touch training, Must initiate training, Punish incorrect training*), followed by two test period (VD task, VD Reversal (VDR) task). Mice were individually housed at least 7 days prior to the start of the experiment (Figure 1a). Food consumption was restricted to maintain mice at 85–90% of their initial free-feeding body weight. Mice were fed every day after the completion of the task session. Food was given immediately after the end of the task, i.e., all mice were removed from the chamber and had access to food as soon as they returned to their home cages. Water was always available in the home cage.

**Figure 1:**
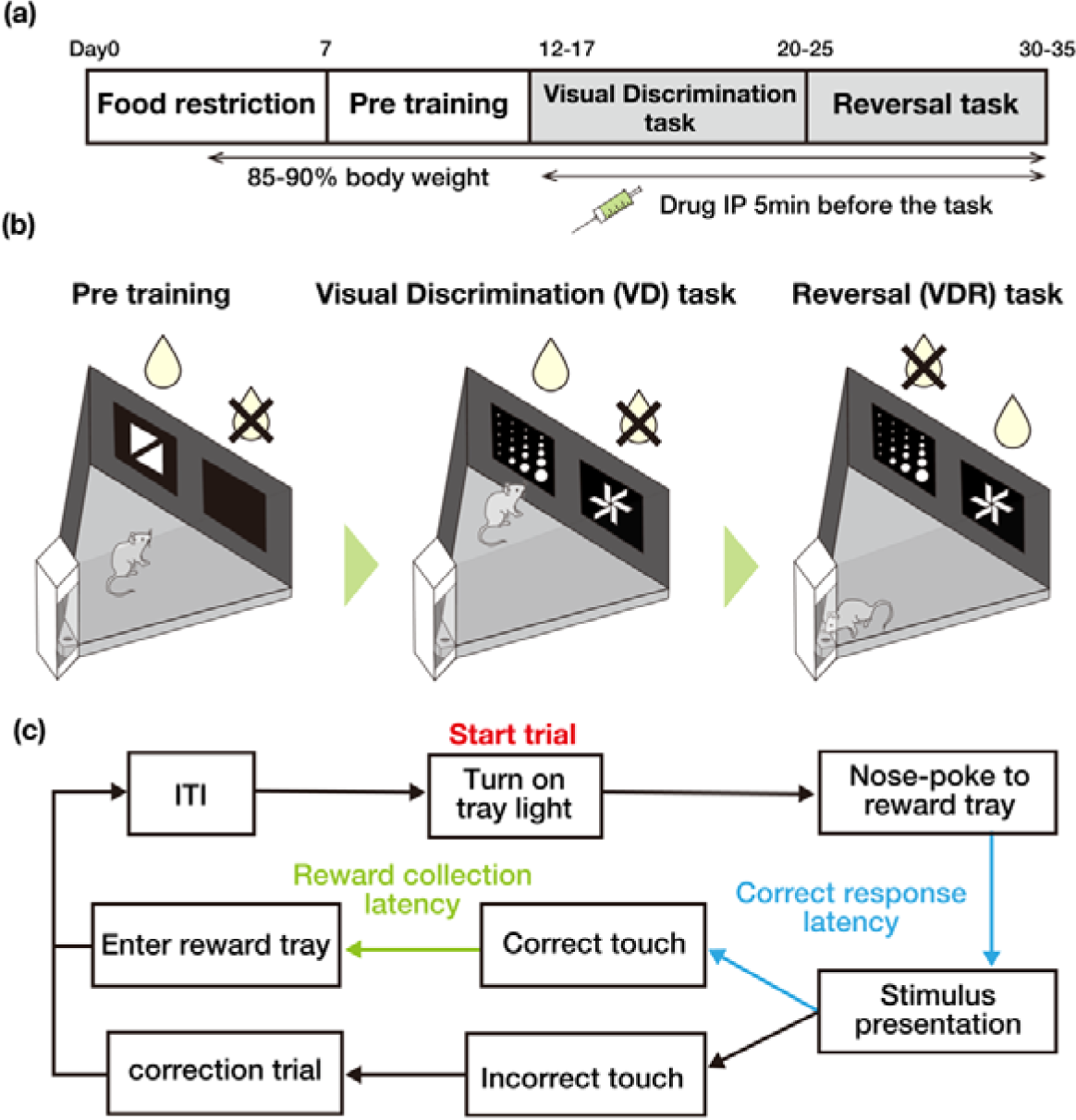
Experimental paradigm. (a) Overall schedule of behavioral experiments. Mice were restricted to 85%∼90% of their body weight at day0 within 1 week. IP: Intraperitoneal administration. (b) The schematic diagram of the Bussey-Saksida touchscreen operant chamber and example visual stimuli in the pre-training and each task. (c) Task design for the Visual Discrimination (VD) task and Reversal task. Correct response latency is the time period indicated by blue arrows and reward collection latency is the period indicated by green arrows.

In *Initial touch training*, during each trial, a visual stimulus was randomly presented on only one panel (Figure 1b). If the mouse did not touch the panel for 30 seconds, a “regular” reward (7μL, 20% condensed milk, Morinaga Milk Industry Co. Ltd., Tokyo, Japan) was delivered. if the mouse touched the panel within 30 seconds, a “tripled” reward (21μL) was delivered immediately. Rewards were delivered with panel lights off, reward tray lights on, and with an auditory cue (3 kHz, 1 second). When the mouse nose-poked into the reward tray to collect the reward, the reward tray light was turned off and a 20-second interval (ITI) period was initiated. At the end of the ITI, the next trial was initiated. The stimulus was displayed no more than four times consecutively in the same position (left or right panel). Sessions were terminated after completion of 30 trials or after 60 minutes. Mice progressed to *Must touch training* following the completion of 30 trials in 60 minutes.

In *Must touch training*, rewards (7 μL) were delivered only when the mouse touched the visual stimulus panel. Other conditions were the same as in *Initial touch training*. Mice progressed to *Must initiate training* following the completion of 30 trials in 60 minutes.

In *Must initiate training*, an additional requirement was added, where the mouse was required to make a nose-poke entry into the reward tray to initiate the trial after the completion of a 20-second ITI. Other conditions were the same as in *Must touch training*. Mice progressed to *Punish incorrect training* following the completion of 30 trials in 60 minutes.

In the *Punish incorrect training*, when the mice were exposed to the incorrect stimulus, the house light was turned on for 5 seconds to indicate the wrong response. After the house lights were turned off and 20 seconds of ITI, a correction trial (image shape and left-right position were the same as in the previous trial) was initiated, and the correction trial was repeated until the mouse touched the correct stimulus. Other conditions were the same as in *Must initiate training*. Mice progressed to the VD task following completion of 30 trials in 30 minutes and achieved at least 23/30 correct responses (76.7 %) on two consecutive days (excluding the number of correction trials).

### 3.5 Visual discrimination (VD) task

Two different visual stimuli, a dot array pattern and a star pattern, were assigned as either the correct or the incorrect stimulus in a counterbalanced manner among mice and kept consistent throughout VD sessions (Figure 1b). The position (left or right panel) of the correct and incorrect visual stimuli were randomized across trials, with the same stimulus displayed no more than four times consecutively in the same position. Other conditions were the same as in *Punish incorrect training* (Figure 1c). The criterion was defined as the completion of 30 trials in 60 minutes and achievement of at least 24/30 correct answers (80.0 %) on two consecutive days. In order to compare learning curves between all individuals, all mice performed the task for 8 days before progressing to the Reversal task, regardless of how fast they reached the criterion. Nicotine at different doses (0.125, 0.25, and 0.5 mg/kg), or saline as a control, were administered via intraperitoneal (i.p.) injection to separate experimental groups 5 minutes prior to each VD session. One female was excluded from the data because it had health problems before the end of the VD task period, and the experiment was terminated.

### 3.6 Reversal (VDR) task

In the VDR task, the correct and incorrect stimuli from the previous stage (VD task) were inverted (i.e., if the star pattern stimulus had been correct during the VD task, the dot array pattern stimulus was now correct during the VDR task, and vice versa for the incorrect stimulus) (Figure 1b). Other conditions were the same as in the VD task. The criterion was defined as the completion of 30 trials in 60 minutes and achievement of at least 24/30 correct answers (80.0 %) on two consecutive days. All mice performed this task for 10 days regardless of how fast they reached the criterion. As with the previous stage, 5 minutes prior to each VDR session, nicotine at different doses (0.125, 0.25, and 0.5 mg/kg), or saline as a control, were administered via i.p. injection to the same experimental groups as had received them during the VD task. One male was excluded from the data because it had health problems before the end of the VDR task period, and the experiment was terminated.

### 3.7 Tissue sampling

Male and female mice housed under identical conditions to mice used for behavioral experiments were administered daily with a 0.5 mg/kg dose of nicotine or saline as a control. After 3 and or 18 days, mice were dissected 65 minutes after nicotine or saline administration. After deep anesthesia with isoflurane and cervical dislocation, brains were removed and immediately cooled and washed in 4°C saline. Brains were sliced into 1-mm thick coronal sections using a brain matrix (Brain Science Idea, Osaka, Japan). The prefrontal cortex (PFC) (Centered on AP = 2.5, ML = 0, DV = -2, mm based on bregma.) was harvested with a 2-mm biopsy punch, immediately placed in 200 μl of RNAlater (Thermo Fisher Scientific, Massachusetts, USA), and stored at 4°C for 24 hours, after which time excess RNAlater solution was removed and samples were frozen at -80°C. Three samples were prepared for each condition.

### 3.8 Transcriptome analysis (RNAseq)

RNA was extracted from cells using a RNeasy plus mini kit (QIAGEN, Venlo, The Netherlands) according to the manufacturer’s protocol. Library preparation for RNA sequencing (RNAseq) was performed using a TruSeq stranded mRNA sample prep kit (Illumina, California, USA) according to the manufacturer’s instructions. Sequencing was performed on an Illumina NovaSeq 6000 sequencer (Illumina) in the 101-base single-end mode. Sequenced reads were mapped to the mouse reference genome sequences (NCBI-RefSeq-GCF_000001635.27_GRCm39) using STAR-2.7.11b. The fragments per kilobase of exon per million mapped fragments (FPKMs) was calculated using RSEM-1.3.3 and TCC-GUI (Su et al., 2019). In TCC-GUI, DEseq2 was used to normalize counts and detect differential expression amongst genes (p<0.05, |FC|<1.1). Pathway analysis was performed using Enrichr (E. Y. Chen et al., 2013; Kuleshov et al., 2016; Xie et al., 2021). Protein-protein interaction (P-PI) networks and cluster enrichment was analyzed using STRING version 12.0 (von Mering et al., 2003, 2005; Szklarczyk et al., 2023). Transcription Factor (TF) analysis was performed using wPGSA (Kawakami et al., 2016). All results are collected in the supplementary tables (TCC-GUI: TableS2, Enrichr pathway analysis: TableS3, STRING cluster enrichment analysis: TableS4, wPGSA TF analysis: TableS5).

### 3.9 Statistical analysis

GraphPad Prism software (v.10.2.0, GraphPad Software Inc., California, USA) was used for all statistical analyses. A repeated-measures two-way ANOVA was used for analyzing the percentage of correct responses, the correct response latency, and the reward collection latency. A one-way ANOVA was used to analyze sessions-to-criterion. For all ANOVAs, post-hoc Dunnett’s multiple comparisons tests were used for comparisons between nicotine and saline groups. All results of ANOVA are collected in the supplementary table (TableS1).

## 4 Results

### 4.1 Nicotine impairs discrimination learning in female, but not male, mice

To examine the effects of nicotine administration on discrimination learning, an important cognitive skill for decision-making, we evaluated task performance of mice in the VD task. In each session, mice were administered nicotine or saline 5 minutes before the start of the task. Male mice administered saline or low (0.125 mg/kg), medium (0.25 mg/kg), or high (0.5 mg/kg) doses of nicotine did not significantly differ in the percentage of correct responses in the VD task (Dose, F(3, 288) = 0.1402, P = 0.9358□; Dose x Days, F(21, 288) = 0.9014, P = 0.5896□; Figure 2a), or in the number of sessions taken to reach the criterion (Dose; F = 0.5833, P = 0.6299□; Figure 2b). Similarly, no significant difference was found between groups in the correct response latency, a measure of locomotion, and in the reward collection latency, a measure of the motivation for the food reward (Supplementary Figure S1).

**Figure 2:**
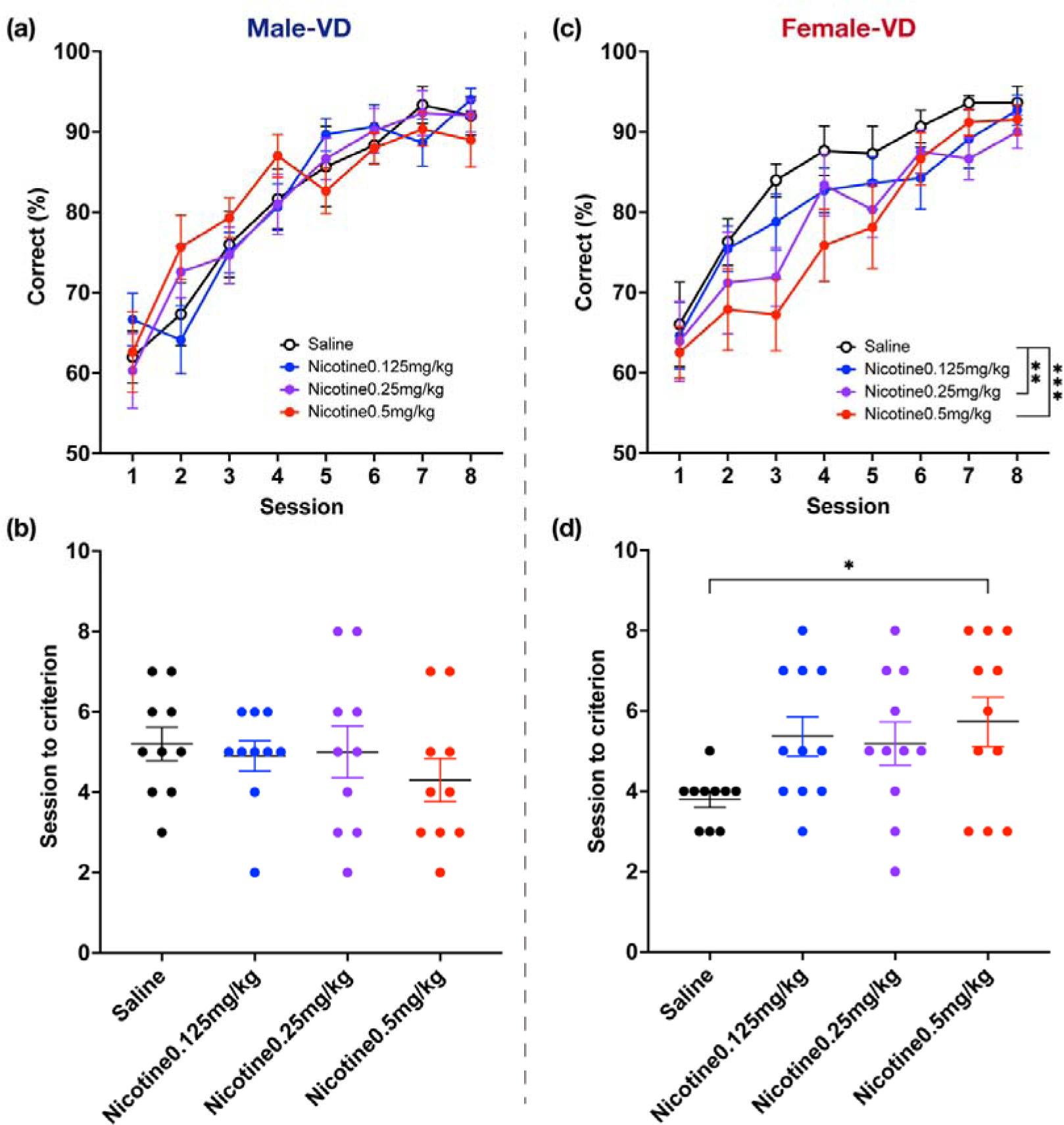
Performance on the VD task. Percentage of correct responses in each session and the number of sessions required to reach the criterion in male (a-b) and female (c-d) mice. Data represent the mean ± SEM, *p < 0.05, **p < 0.01, ***p < 0.001 in post-hoc analyses after two-way ANOVA and one-way ANOVA (Male; Saline n=10, Nicotine 0.125 mg/kg n= 10, Nicotine 0.25 mg/kg n= 10, Nicotine 0.5 mg/kg n= 10, Female; Saline n=10, Nicotine 0.125 mg/kg n= 11, Nicotine 0.25 mg/kg n= 11, Nicotine 0.5 mg/kg n= 11).

In contrast, in female mice, medium and high doses, but not a low dose, of nicotine significantly decreased the percentage of correct responses compared to those administered saline (Dose, F(3, 312) = 5.992, P = 0.0006□; Dose x Days, F(21, 312) = 0.5884, P = 0.9256□; Figure 2c). In addition, high-dose nicotine significantly increased the number of sessions to reach the criterion (Dose, F = 2.744, P = 0.056; post hoc, saline vs high dose, AdjP = 0.0269; Figure 2d). Similar to male mice, no significant differences were found between groups in the correct response latency and the reward collection latency (Supplementary Figure S1).

These findings indicate that nicotine impairs VD learning in female mice without affecting locomotion or motivation.

### 4.2 Nicotine increases cognitive flexibility in male mice but decreases it in female mice

Next to explore the effect of nicotine on cognitive flexibility, the ability to adaptively shift mental strategies based upon the changing demands of the environment, mice were tested in the VDR task. In this test of reversal learning, the correct and incorrect images from the previous VD task were reversed.

In male mice, high-dose nicotine significantly increased the percentage of correct responses when compared with saline (Dose, F(3, 230) = 4.579, P = 0.0039; Dose x Days, F(27, 230) = 0.4542, P = 0.9916; Figure 3a), while low and medium doses of nicotine had no significant effect. Additionally, high-dose nicotine significantly decreased the number of sessions to reach the criterion when compared with saline (Dose, F = 2.432, P = 0.0909; post hoc, saline vs high dose, AdjP = 0.0378; Figure 3b). No significant differences were found between groups in the correct response latency and the reward collection latency (Supplementary Figure S2).

**Figure 3:**
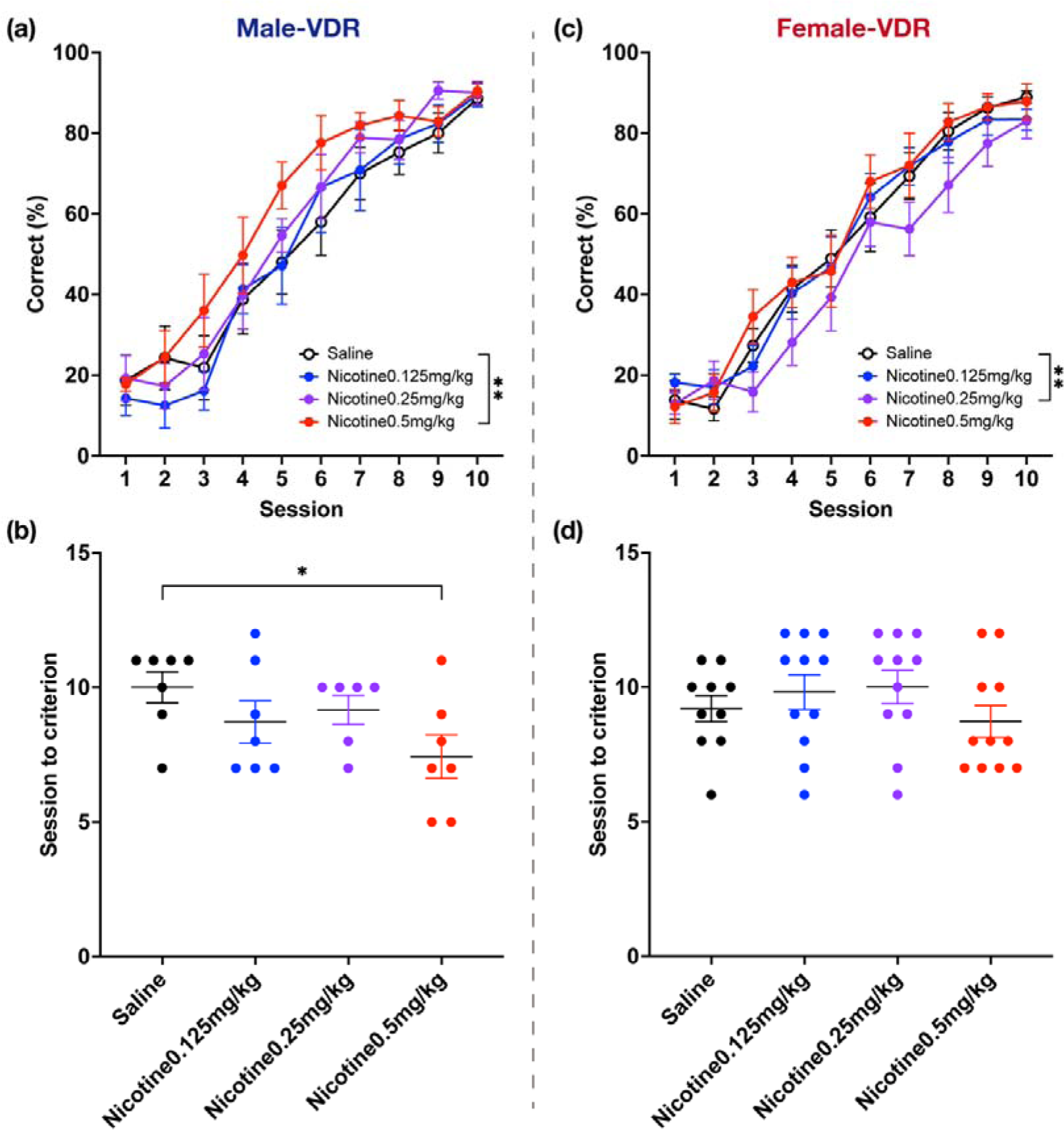
Performance on the reversal (VDR) task. Percentage of correct responses in each session and the number of sessions required to reach the criterion in male (a-b) and female (c-d) mice. Data represent the mean ± SEM, **p < 0.01 in post-hoc analyses after two-way ANOVA and one-way ANOVA (Male; Saline n=10, Nicotine 0.125 mg/kg n= 10, Nicotine 0.25 mg/kg n= 10, Nicotine 0.5 mg/kg n= 10, Female; Saline n=10, Nicotine 0.125 mg/kg n= 11, Nicotine 0.25 mg/kg n= 11, Nicotine 0.5 mg/kg n= 11).

In contrast, in female mice, medium-dose nicotine significantly decreased the percentage of correct responses when compared with saline, while low and high doses nicotine dose had no significant effect (Dose, F(3, 390) = 5.321, P = 0.0013; Dose x Days, F(27, 390) = 0.5004, P = 0.9840□; Figure 3c). No significant effect of nicotine dose was observed on sessions to reach the criterion (Dose; F = 0.9865, P = 0.4091; Figure 3d). Similarly, no significant differences were found between groups for the correct response latency and the reward collection latency (Supplementary Figure S2).

Taken together, these results reveal that high-dose nicotine administration increases cognitive flexibility in a VDR task in male mice, while medium-dose nicotine decrease cognitive flexibility in female mice.

### 4.3 Genes upregulated after 3 days of nicotine administration

To elucidate cellular alterations resulting from repeated nicotine administration, as well as their differences in males and females, RNAseq was performed on brain tissue from nicotine-treated mice. Two different time points were selected for RNAseq experiments; after 3 days of nicotine administration, as a measure of short-term nicotine administration (and a time point that was sufficient to induce cognitive differences in VD performance in female mice), and after 18 days of nicotine administration, as a measure of long-term nicotine administration (and a time point that represents the most amount of administrations that mice were exposed to in behavioral experiments). As the high-dose (0.5.mg/kg) of nicotine produced the greatest changes in performance in VD and VDR tasks, this dose was chosen for RNAseq experiments. Finally, the PFC was chosen as a target for analysis as this brain region has previously been implicated in several cognitive abilities, including discrimination learning and cognitive flexibility (Friedman & Robbins, 2022). Thus, differential gene expression in the PFC was analyzed in four groups, on days 3 and 18 in both males and females. Subsequently, using gene sets with altered expression, P-PI network analysis, pathway analysis, and TF analysis were performed.

P-PI network analysis on the Day3-male upregulated gene set generated one large network of six clusters and four small networks. The largest cluster identified by cluster (cluster-by-cluster) enrichment analysis included genes related to “oligodendrocyte differentiation”, “axon sheath”, and “myelin”. The second largest cluster was related to the “PI3K-Akt-mTOR signaling pathway” (Figure 4a, Table S3). No terms were significantly enriched in the pathway analysis when using the two databases (Figure 4b,c). In Day3-female mice, P-PI network analysis generated one large network of three clusters and six smaller networks. In cluster enrichment analysis, “Collagen-containing extracellular matrix” and “PI3K-Akt-mTOR signaling pathway” were enriched in the components of the largest cluster component, “AP-1 transcription factor” and “MAP kinase activation” in the second cluster, and “MHC class □ protein complex” in the third cluster (Figure 4d, Table S3). Pathway analysis showed significant enrichment of “Oncostatin M”, “Beta-1 integrin cell surface interaction”, “BDNF signaling pathway”, “Integrins in angiogenesis”, and “FRA pathway” in the BioPlanet2019 database, as well as “Integrins in angiogenesis”, and “FRA pathway” in Reactome2022 (Figure 4e,f). The female gene set had more significantly enriched terms compared to the male set.

**Figure 4:**
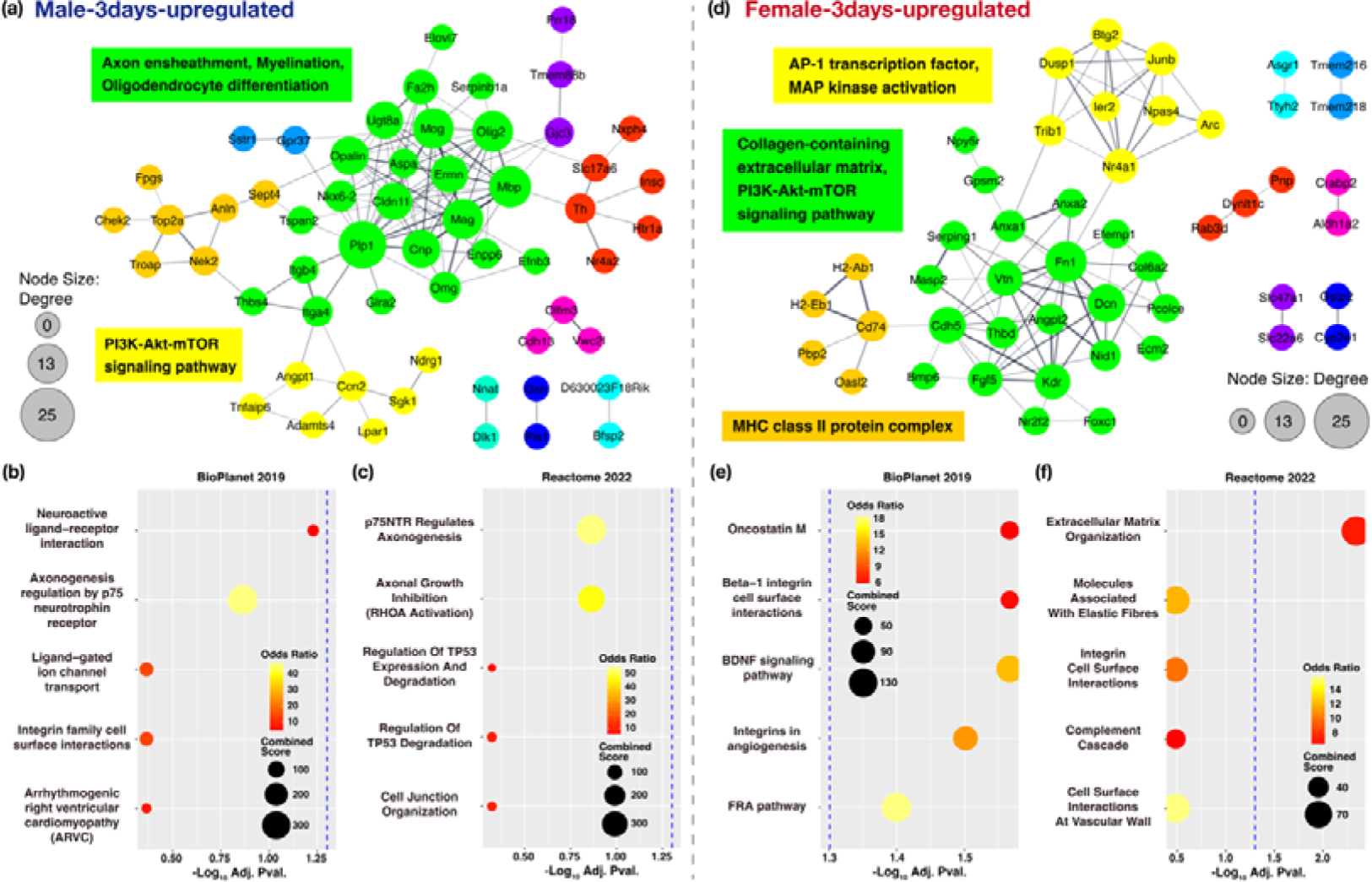
Analysis of the 3days-upregureted gene set. Results of the P-PI network cluster analysis and pathway analysis in male (a-c) and female (d-f) mice. Cluster and enriched terms for the largest (green), second (yellow), and third (orange) clusters of the P-PI network. Blue dashed line indicates p=0.05 in the pathway analysis. The results of the cluster enrichment and pathway analyses are summarized in the supplementally table (Table S3-4).

### 4.4 Genes upregulated after 18 days of nicotine administration

Using the Day18-male upregulated gene set, P-PI network analysis generated two large networks of four clusters and fourteen small networks. Components of the largest clusters were genes related to “AP-1 transcription factor” and “MAP kinase activation” by cluster enrichment analysis (Figure 5a, S3). No significant terms were enriched in pathway analysis (Figure 5b,c). In Day18-female mice, the P-PI network analysis generated one large network of twelve clusters and two smaller networks. In cluster enrichment analysis, “Integrin binding”, “Growth factor activity”, “PI3K-Akt signaling pathway”, and “TGF-beta signaling pathway” were enriched in components of the largest cluster, “Cell Cycle”, “Checkpoints”, and “Mitotic” in the second cluster, “MHC class □ protein complex” in the third cluster, “Innate immune response” in the fourth cluster, and “Myelination” in the fifth cluster (Figure 5d, Table S3). In addition, a small cluster of Fos and Junb, which are AP-1 transcription factors, was found to interact with 6 clusters. Pathway analysis showed significant enrichment of “Interleukin-4 regulation of apoptosis”, “BDNF signaling pathway”, “Neural crest differentiation”, “Transport of glucose and other sugars”, “bile salts and organic acids”, “metal ions and amine compounds”, and “TGF-beta regulation of extracellular matrix” in BioPlanet 2019 (Figure 5e,f). Across all upregulated gene sets, there were more significantly enriched terms in female sets than in the male sets, and an increase of enriched terms as a result of long-term administration was seen only in females.

**Figure 5:**
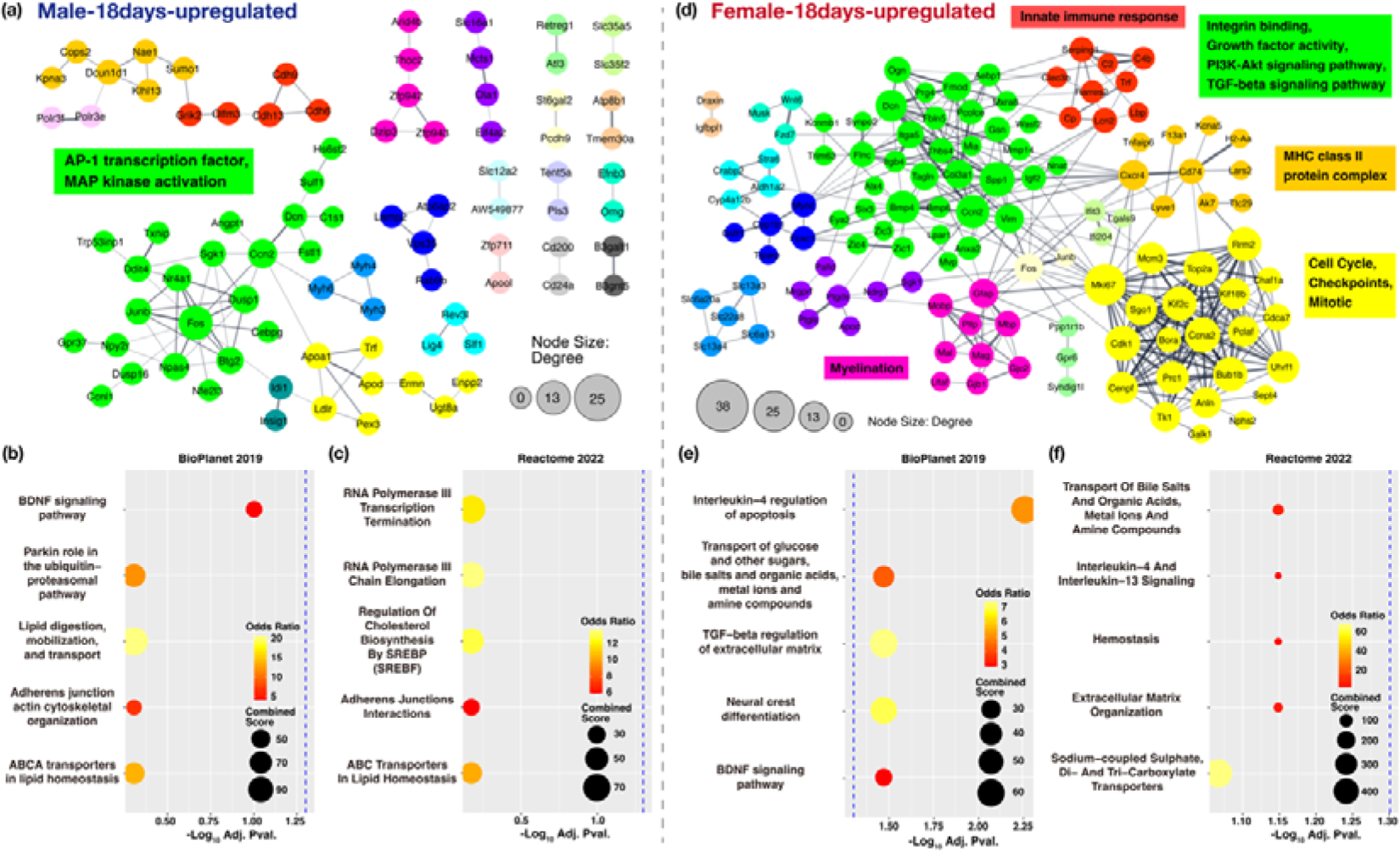
Analysis of the 18days-upregureted gene set. Results of the P-PI network cluster analysis and pathway analysis in male (a-c) and female (d-f) mice. Cluster and enriched terms for the largest (green), second (yellow), third (orange), fourth (red), and fifth (pink) clusters of the P-PI network. Blue dashed line indicates p=0.05 in the pathway analysis. The results of the cluster enrichment and pathway analyses are summarized in the supplementally table (Table S3-4).

### 4.5 Genes downregulated after 3 days of nicotine administration

Using the Day3-male downregulated gene set, P-PI network analysis generated one large network of three clusters and eight small networks. Components of the largest clusters were genes related to “Glutamatergic synapse and Postsynaptic density”, and the second cluster included genes related to “Transmembrane receptor protein tyrosine kinase signaling pathway” (Figure 6a, Table S3). “Neural System” was only significantly enriched in Reactome2022 (Figure 6b,c). In Day3-female mice, the P-PI network analysis generated five small networks (Figure 6d, Table S3), and no significant terms were enriched in the pathway analysis (Figure 6e,f). These findings indicate that nicotine has no significant suppressing effect on gene expression in female mice.

**Figure 6:**
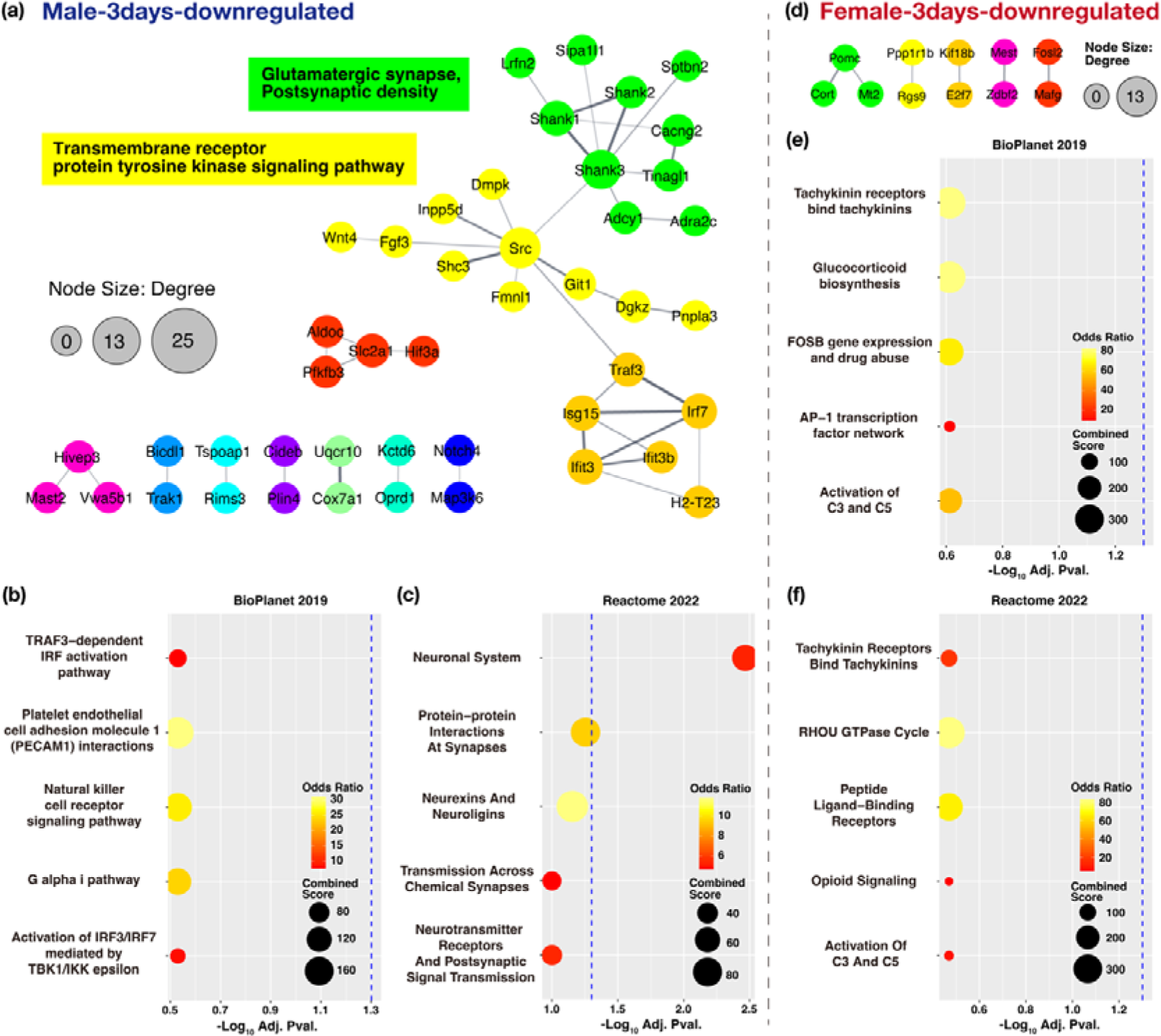
Analysis of the 3days-downregureted gene set. Results of the P-PI network cluster analysis and pathway analysis in male (a-c) and female (d-f) mice. Clusters and enriched terms for the largest (green) and second (yellow) clusters of the P-PI network. Blue dashed line indicates p=0.05 in the pathway analysis. The results of the cluster enrichment and pathway analyses are summarized in the supplementally table (Table S3-4).

### 4.6 Genes downregulated after 18 days of nicotine administration

Using the Day18-male downregulated gene set, P-PI network analysis generated one large network of nineteen clusters and twelve small networks. In cluster enrichment analysis, “Glutamatergic synapse”, “Neurexins and neuroligins”, and “Cell junction”, “Postsynapse”, were enriched in components of the largest cluster, “Transmembrane receptor protein tyrosine kinase signaling pathway” and “Focal adhesion” in the second cluster, and “Histone deacetylase binding” and “Negative regulation of transcription by RNA polymerase II” in the third cluster (Figure 7a, Table S3). Pathway analysis revealed several pathways that were significantly enriched, including “Neuronal system”, “L1CAM interactions”, “Interaction between L1-type proteins and ankyrins”, “PGC-1a regulation”, “Endocytosis”, “Neurexins and Neuroligins”, “Protein-protein Interactions at Synapses”, “Transmission Across Chemical Synapses”, and “Activation of NMDA Receptors and Postsynaptic Events” (Figure 7b,c). In female mice, P-PI network analysis generated only one small network (Figure 7d, Table S3), And no significant terms were enriched in the pathway analysis (Figure 7e,f). Therefore, regardless of the number of doses, nicotine has very little suppressing effect on gene expression in female mice. On the other hand, far more enriched terms were found in the male gene set, and the size of the networks and the number of enriched terms increased as a result of long-term nicotine administration.

**Figure 7:**
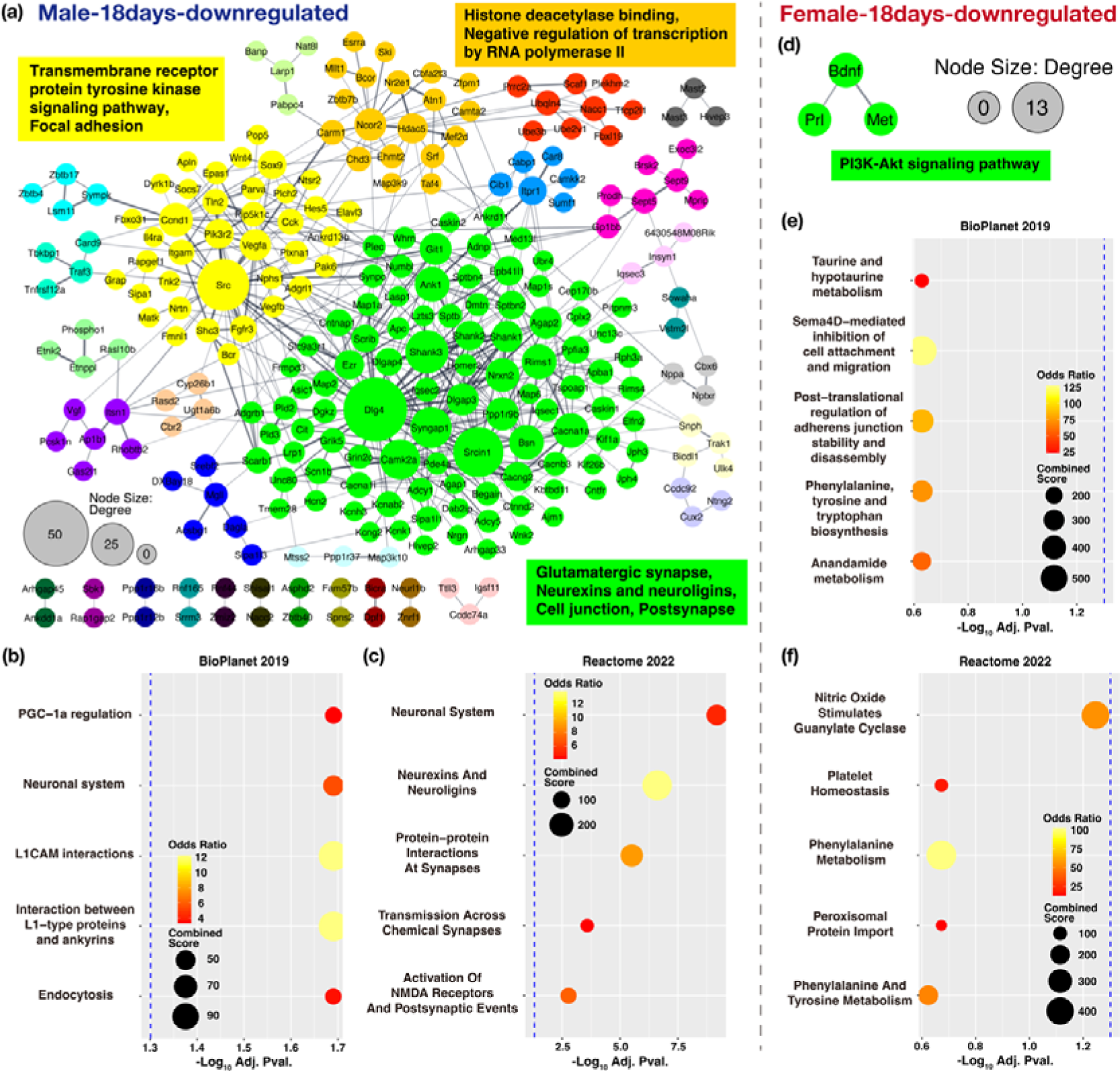
Analysis of the 18days-downregureted gene set. Results of the P-PI network cluster analysis and pathway analysis in male (a-c) and female (d-f) mice. Clusters and enriched terms for the largest (green), second (yellow), and third (orange) clusters of the P-PI network. Blue dashed line indicates p=0.05 in the pathway analysis. The results of the cluster enrichment and pathway analyses are summarized in the supplementally table (Table S3-4).

## 5 Discussion

Here, for the first time, we evaluated the sex-dependent effects of nicotine on discrimination learning and cognitive flexibility in mice. In male mice, nicotine significantly increased performance in the VDR task, while in female mice, nicotine significantly impaired performance in the VD task.

Our findings contrast somewhat with some previous studies in male mice, which reported chronic nicotine administration with osmotic pumps to enhance VD performance, but inhibit VDR performance (Cole et al., 2015; Ortega et al., 2013). However, other groups have reported that acute nicotine in male rats is able to facilitate reversal learning in a probabilistic task (Milienne-Petiot et al., 2017). The reason for the opposing findings of studies into the effects of nicotine on cognition is still unclear, but may be due to differences in the method (i.p. injection, oral, osmotic pump) and duration (acute vs chronic) of nicotine administration. Additionally, even when the same task is used, different designs may result in different aspects of cognition being measured. For example, VD and VDR tasks in previous studies have largely been performed in “classic” operant chambers where the visual cue is a single cue light illuminated above the correct or incorrect lever (Cole et al., 2015; Ortega et al., 2013; Milienne-Petiot et al., 2018). Thus, mice/rats are required only to attend to a single cue and to learn to approach or avoid that cue. However, in our touchscreen chamber tasks, correct and incorrect visual cues are presented simultaneously, likely requiring greater attentional control to discriminate between the visual patterns in order to make the correct decision.

Interestingly, in our experiments, nicotine consistently impaired performance in the VD task in female mice, suggesting that the previously reported beneficial effects of nicotine on performance may be limited to male mice and may have opposite effects in females. Indeed, a previous study in rats reported nicotine to dose-dependently increase impulsivity in females, but to only increase impulsivity in males at a high dose. It is possible that in the present study, nicotine administration may have caused females to impulsively respond at the previously correct response window during the VDR task, explaining their worsened performance. This notion is supported by evidence that female rats are known to express higher levels of nAChRs (Koylu et al., 1997), whose expression in the medial PFC has been linked with impulsivity (Ohmura et al., 2012). Additionally, nicotine is reported to accumulate faster, distribute over a larger area, and be metabolized slower in female rats than in males (Rosecrans, 1972).

In previous studies, acute and chronic nicotine increased the motivation for sucrose reward in rats (Grimm et al., 2012; Jias & Ellison, 1990; Lacy et al., 2012). On the other hand, acute and self-administration of nicotine has been reported to decrease the motivation or intake of sucrose in rats (Bunney et al., 2016; Hart et al., 2021). Similarly, smoking was reported to decrease food intake in mice (H. Chen et al., 2005); however, nicotine pre-exposure did not influence the reinforcing effects of sucrose in rats (Schwartz et al., 2018). Thus, the effects of nicotine on motivation for food rewards are inconsistent. Analysis of the reward collection latency in our VD and VDR experiments revealed no changes in motivation for food rewards following nicotine administration, supporting the previous findings ofSchwartz et al. (2018). This data also suggests that nicotine-induced changes in performance in VD and VDR tasks in the present study are not attributable to alterations in motivation.

Next, to explore how nicotine may differentially alter gene expression between the sexes, we performed transcriptomic analysis of the PFC, a brain region linked with discrimination learning and cognitive flexibility, in male and female mice administered with nicotine over short (3 days) or long (18 days) durations. Our analyses revealed three categories of cellular alterations induced by nicotine administration. The first category involved common alterations shared between both sexes, with the most notable effect observed in TF activity. Approximately one-third of the altered TFs were common across all groups (day and sex) (Supplementary Figure S3). These common changes in TFs likely reflect the acute effects of nicotine, which occur with every administration regardless of sex. TFs that were altered across all groups included the Myc/Max/Mad network components that regulate histone acetylation and several demethylases. Furthermore, cluster enrichment analysis on the P-PI network showed that nicotine increased the expression of PI3K-Akt signaling pathway-related genes and/or ΑP-1 transcription factors such as Fos and Junb, regardless of day and sex. Nicotine is known to activate the PI3K-Akt signaling pathway via nAChRs, through which it modulates glucose metabolism, cell cycle progression, and apoptosis (He et al., 2024; West et al., 2003). Nicotine is also known to induce the expression of immediate early genes such as c-fos and junB in various brain regions (Emilio Merlo Pich et al., 1997; Nisell et al., 1997; Schilström et al., 2000), and our results are consistent with these findings. Thus, these are most likely early cellular responses to nicotine across male and female mice. Previous qPCR array experiments have shown that gestational nicotine exposure upregulates expression of the major myelin genes in male mice and downregulates it in female mice (Cao et al., 2013). The first cluster of the male-3days-upregulated analysis contained the major myelin genes, consistent with previous studies; however, the fifth cluster of the female-18days-upregulated analysis also contained the major myelin genes, which is in the opposite direction to previous results. This may indicate that, at least for myelin genes, the effects of nicotine on gene expression may be reversed between embryonic and adult stages.

The second category involved male-specific effects of nicotine seen in the downregulated gene set. P-PI network cluster analysis showed that “glutamatergic synapses”, “post synapse”, and “neurexins and neuroligins” were terms enriched in the largest cluster centered on the Shank family and Dlg4 (PSD-95). Additionally, the “transmembrane receptor protein tyrosine kinase signaling pathway” term was enriched in the second cluster centered on the Src gene, regardless of the number of doses. The “neuronal system” term was also significantly enriched in the pathway analysis. Interestingly, nicotine-induced downregulation of gene expression appears to become stronger as the number of administrations increases, as the cluster size was larger in the 18-day treatment compared to the 3-day treatments. Shank family, Dlg4 (PSD-95), and neuroligins are proteins with postsynaptic functions in excitatory synapses and are known to be risk factors for autism (Berkel et al., 2010; Durand et al., 2007; Michael Feyder et al., 2010; Pinto et al., 2010; Sato et al., 2012; Südhof, 2008). Mice deficient in Shank, Neuroligin, or PSD-95 complex components have previously been evaluated as mouse models of autism (Qin et al., 2018; Radyushkin et al., 2009; Winkler et al., 2018). SHANK3Δ11 deficiency is known to reduce cognitive flexibility, and Shank3B heterozygous mice demonstrate impaired discrimination learning (Copping et al., 2017; Ponzoni et al., 2019). Among the components of the PSD-95 protein complex, Syngap1 heterozygous mice have been reported to show impaired cognitive flexibility, whereas deletion of neuroligin-3 or Dlgap2 enhanced cognitive flexibility (Horner et al., 2021). In addition, mice with autism-associated mutations in neuroligin-3 showed improved attention to stimuli on the rodent continuous-performance test at low difficulty levels (Burrows et al., 2022). Thus, postsynaptic components have been shown to alter cognitive flexibility, and our findings that nicotine administration downregulated postsynaptic components may help to explain the increased cognitive flexibility in male mice administered nicotine.

The final category involved the female-specific effects of nicotine. In the upregulated gene set at both time points (3days and 18days), P-PI network cluster analysis showed enrichment of the MHC (major histocompatibility complex) class II protein complex. The MHC class II protein complex is known to be constitutively expressed in antigen-presenting cells, and intracellular MHC class II molecules act as adaptors and facilitate full activation of TLR-induced adaptive immunity (Liu et al., 2011; Wieczorek et al., 2017). In addition, the fourth cluster of the P-PI network in Day18-female mice showed enrichment of “innate immune response”. In previous studies with female mice, nicotine has been shown to increase the expression of proinflammatory cytokines (Kumar et al., 2024), which are known to result in upregulation of microglial MHC class II protein expression (Imamura et al., 1994; Neumann et al., 1996). This suggests that nicotine may function as a proinflammatory regulator only in female mice, and that chronic nicotine exposure enhances this effect. On the other hand, there were few downregulated genes, and a highly underdeveloped P-PI network generated from the downregulated gene set compared to males, indicating that nicotine has little effect on gene suppression in female mice. In addition, TF analysis revealed alteration of several additional components of the Myc/Max/Mad network in female mice in addition to the common components altered in both sexes, possibly suggesting that nicotine-induced deacetylation of the Myc/Max/Mad network is stronger in female mice. Innate immune responses can lead to synaptic and circuit dysfunction, which have been shown to increase vulnerability to cognitive decline and neurodegeneration in humans (Cribbs et al., 2012; Haroon et al., 2017; Turner et al., 2021). Thus, it is possible that nicotine inhibits VD performance via activation of innate immune responses in female mice.

In summary, in our studies, nicotine administration was found to facilitate discrimination in male mice, but impair cognitive flexibility in female mice. In our transcriptome analysis, we found a common effect of nicotine in males and females increasing the expression of signaling factors immediately downstream of the nAChRs. On the other hand, we also found sex-dependent effects, particularly reduced expression of postsynaptic-related genes in males and increased expression of innate immunity-related genes in females. Gene set clusters that differed between male and female mice may induce sexually differential effects of nicotine on VD and VDR performance, and further investigation on such intracellular pathways is necessary to further elucidate the molecular differences underlying the sex-dependent differences in the effects of nicotine on cognitive functions.

## Data availability

All sequencing data and gene count table is available on DDBJ BioProject accession ID: PRJDB17923.

## Acknowledgments

The authors thank Ms. Noriko Otani for technical assistance. RNA sequencing was performed at the NGS core facility at the Research Institute for Microbial Diseases of Osaka University.

## Author Contributions

Y.A., Y.S., T.O., and T.H. contributed to the conception and design of the study and interpretation of data. Y.A., Y.S., and M.A. performed the data acquisition and analysis. Y.A., K.S., T.M., T.O., and T.H. wrote the manuscript. All authors contributed to manuscript revision and read and approved the submitted version.

## Abbreviations

nAChRs: Nicotinic acetylcholine receptors
VD task: Visual discrimination
VDR task: Reversal
RNAseq: RNA sequencing
PFC: Prefrontal cortex
P-PI: Protein-protein interaction
TF: Transcription factor

## Supplementary

Other Supplementary Material for this manuscript includes the following: “Supplementary_TableS1-Statistics.xlsx” : Statistics of all behavioral experiments “Supplementary_TableS2-TCC-GUI-DEG-list.xlsx” : Differentially Expressed Genes of TCC-GUI “Supplementary_TableS3-STRING-Cluster-enrichment.xlsx” : Results of cluster enrichment analysis “Supplementary_TableS4-Enricher-Pathway.xlsx” : Results of pathway analysis “Supplementary_TableS5-wPGSA.xlsx” : Results of transcription factor analysis (wPGSA)

**Supplementary Figure S1:**
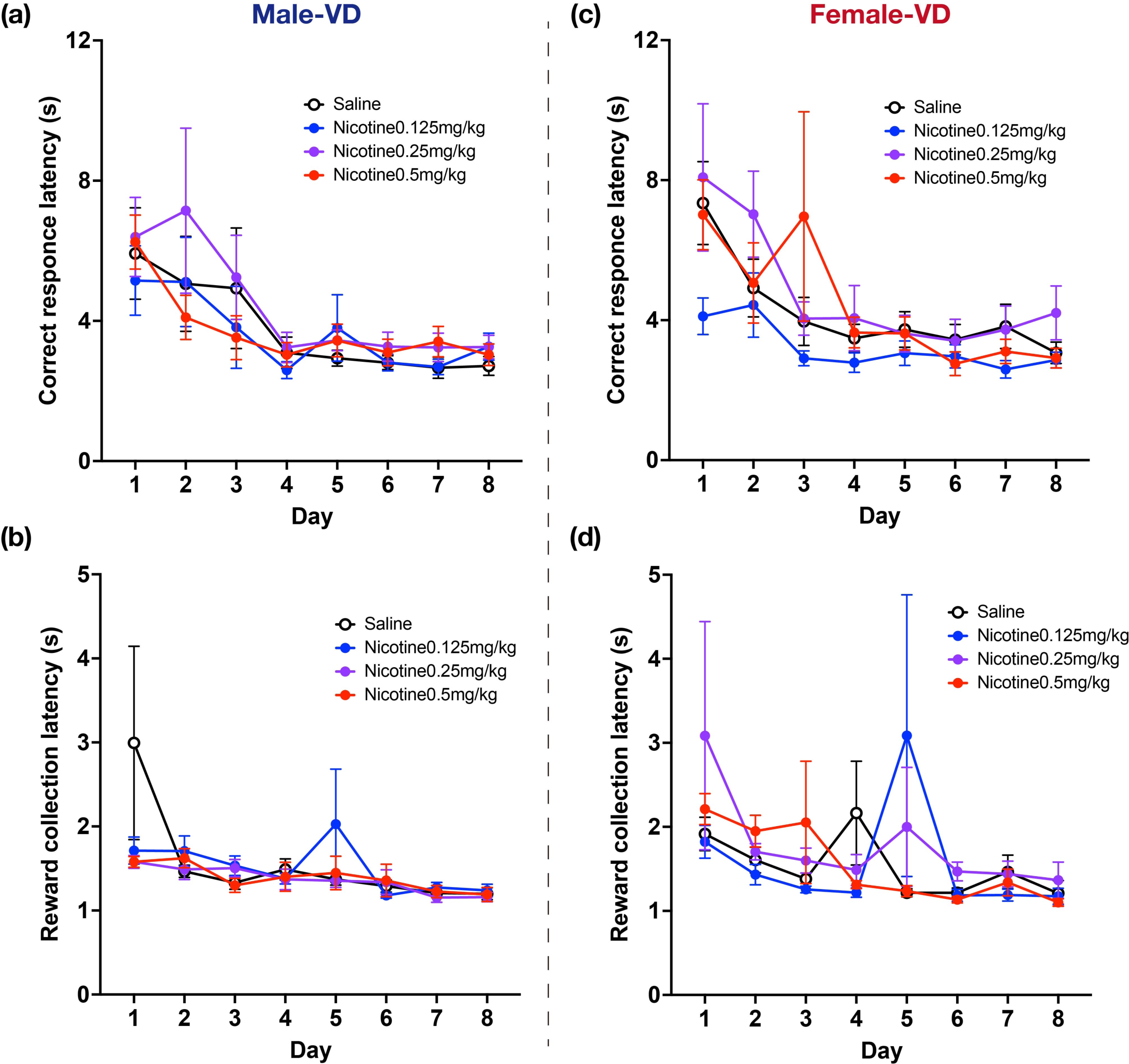
Locomotion and motivation for rewards in the visual discrimination (VD) task. Correct response latency and reward collection latency in each session for male (a-b) and female (c-d) mice. Data represent the mean ± SEM, two-way ANOVA (Male; Saline =10, Nicotine 0.125 mg/kg = 10, Nicotine 0.25 mg/kg = 10, Nicotine 0.5 mg/kg = 10, Female; Saline =10, Nicotine 0.125 mg/kg = 11, Nicotine 0.25 mg/kg = 11, Nicotine 0.5 mg/kg = 11).

**Supplementary Figure S2:**
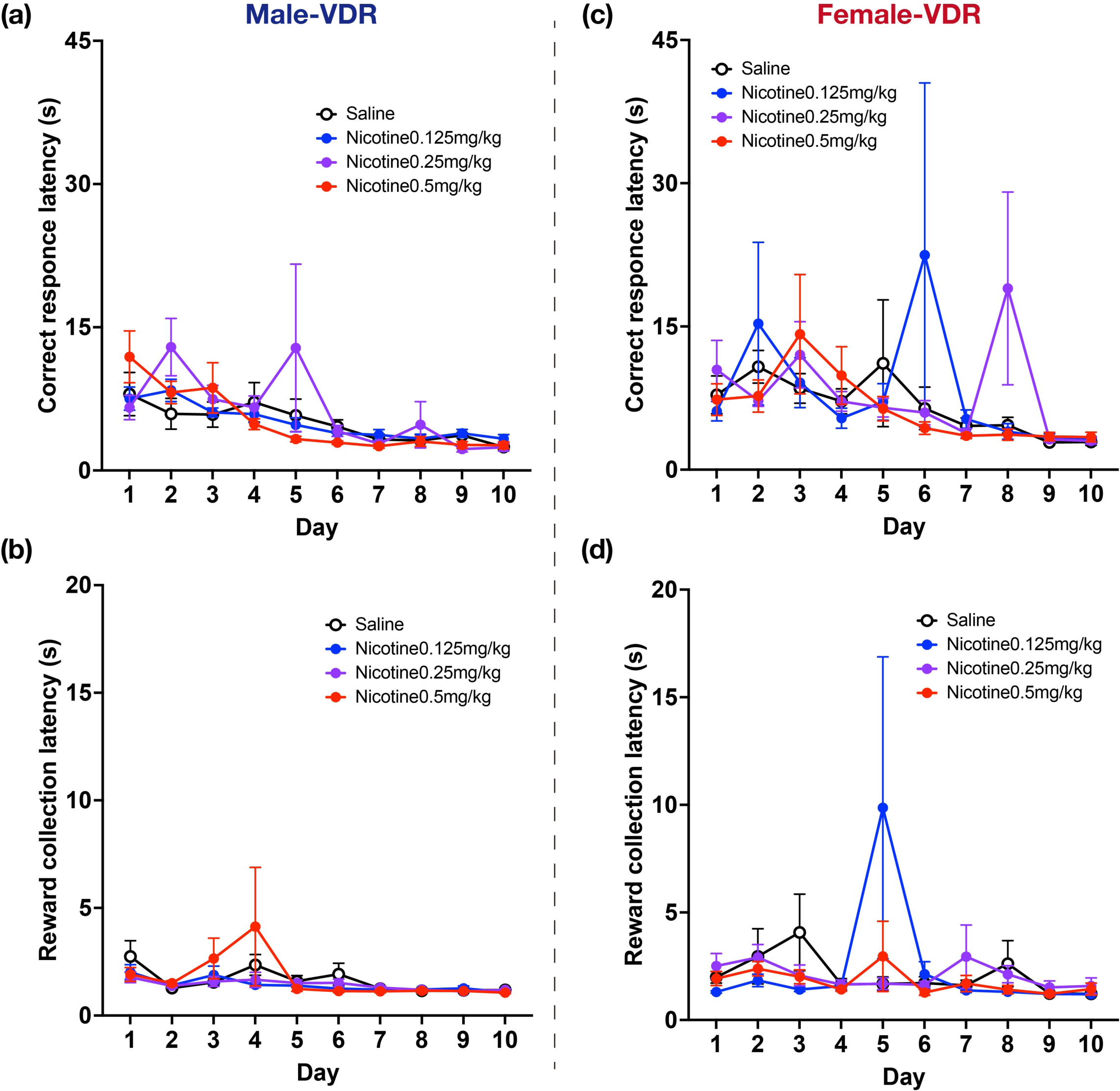
Locomotion and motivation for rewards in the reversal (VDR) task. Correct response latency and reward collection latency in each session for male (a-b) and female (c-d) mice. Data represent the mean ± SEM, two-way ANOVA (Male; Saline =10, Nicotine 0.125 mg/kg = 10, Nicotine 0.25 mg/kg = 10, Nicotine 0.5 mg/kg = 10, Female; Saline =10, Nicotine 0.125 mg/kg = 11, Nicotine 0.25 mg/kg = 11, Nicotine 0.5 mg/kg = 11).

**Supplementary Figure S3:**
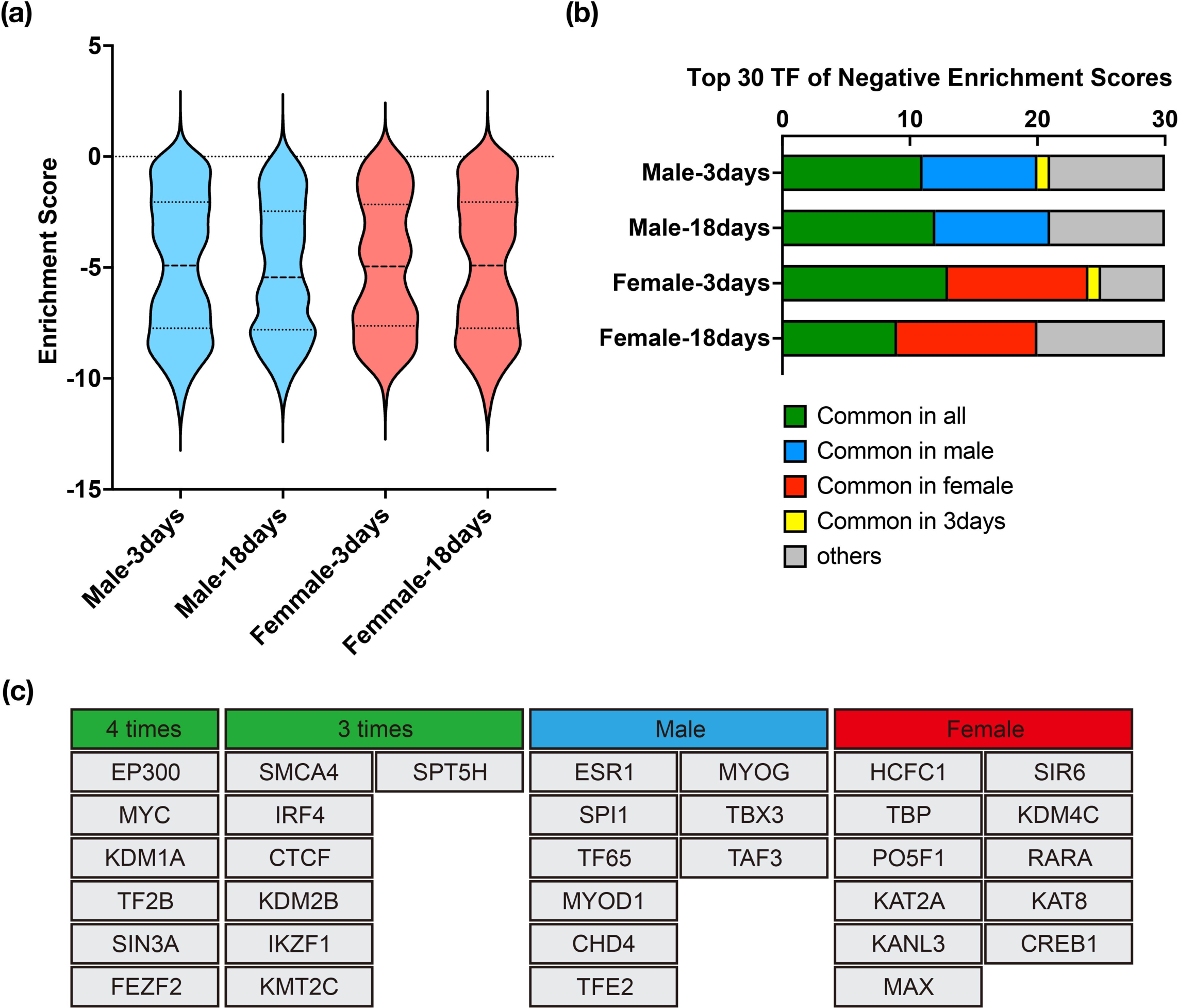
The results of transcription factor (TF) analysis by wPGSA (a) The enrichment score indicates gene expression variation downstream of the TF binding site. 454 TFs were analyzed. (b) The 30 TFs with the lowest enrichment scores in each group and their commonality (c) Individual names of TFs that were common in (b). 4 times and 3 times indicates the number of appearances in all four groups.

## References

Bagdas, D., Diester, C. M., Riley, J., Carper, M., Alkhlaif, Y., AlOmari, D., Alayoubi, H., Poklis, J. L., & Damaj, M. I. (2019). Assessing nicotine dependence using an oral nicotine free-choice paradigm in mice. Neuropharmacology, 157. 10.1016/j.neuropharm.2019.107669

Benowitz, N. L. (2008). Clinical pharmacology of nicotine: Implications for understanding, preventing, and treating tobacco addiction. In Clinical Pharmacology and Therapeutics (Vol. 83, Issue 4, pp. 531–541). 10.1038/clpt.2008.3

Berkel, S., Marshall, C. R., Weiss, B., Howe, J., Roeth, R., Moog, U., Endris, V., Roberts, W., Szatmari, P., Pinto, D., Bonin, M., Riess, A., Engels, H., Sprengel, R., Scherer, S. W., & Rappold, G. A. (2010). Mutations in the SHANK2 synaptic scaffolding gene in autism spectrum disorder and mental retardation. Nature Genetics, 42(6), 489–491. 10.1038/ng.589

Booze, R. M., Welch, M. A., Wood, M. L., Billings, K. A., Apple, S. R., Mactutus, C. F., ¶#§, Booze, R. M., Welch, M. A., Wood, M. L., Billings, K. A., Apple, S. R., & Behav, C. F. M. (1999). Behavioral Sensitization Following Repeated Intravenous Nicotine Administration: Gender Differences and Gonadal Hormones. Pharmacology Biochemistry and Behavior, 64(4), 827–839.

Bunney, P. E., Burroughs, D., Hernandez, C., & LeSage, M. G. (2016). The effects of nicotine self-administration and withdrawal on concurrently available chow and sucrose intake in adult male rats. Physiology and Behavior, 154, 49–59. 10.1016/j.physbeh.2015.11.002

Burrows, E. L., May, C., Hill, T., Churliov, L., Johnson, K. A., & Hannan, A. J. (2022). Mice with an autism-associated R451C mutation in neuroligin-3 show a cautious but accurate response style in touchscreen attention tasks. Genes, Brain and Behavior, 21(1). 10.1111/gbb.12757

Caldarone, B. J., King, S. L., & Picciotto, M. R. (2008). Sex differences in anxiety-like behavior and locomotor activity following chronic nicotine exposure in mice. Neuroscience Letters, 439(2), 187–191. 10.1016/j.neulet.2008.05.023

Cao, J., Wang, J., Dwyer, J. B., Gautier, N. M., Wang, S., Leslie, F. M., & Li, M. D. (2013). Gestational nicotine exposure modifies myelin gene expression in the brains of adolescent rats with sex differences. Translational Psychiatry, 3. 10.1038/tp.2013.21

Chaudhri, N., Caggiula, A. R., Donny, E. C., Booth, S., Gharib, M. A., Craven, L. A., Allen, S. S., Sved, A. F., & Perkins, K. A. (2005). Sex differences in the contribution of nicotine and nonpharmacological stimuli to nicotine self-administration in rats. Psychopharmacology, 180(2), 258–266. 10.1007/s00213-005-2152-3

Cheeta, S., Irvine, E. E., Tucci, S., Sandhu, J., & File, S. E. (2001). In Adolescence, Female Rats Are More Sensitive to the Anxiolytic Effect of Nicotine Than Are Male Rats. Neuropsychopharmacology, 25(4), 601–607. www.acnp.org/citations/Npp

Chen, E. Y., Tan, C. M., Kou, Y., Duan, Q., Wang, Z., Meirelles, G. V., Clark, N. R., & Ma’ayan, A. (2013). Enrichr: interactive and collaborative HTML5 gene list enrichment analysis tool. BMC Bioinformatics, 14(128). 10.1186/1471-2105-14-128

Chen, H., Vlahos, R., Bozinovski, S., Jones, J., Anderson, G. P., & Morris, M. J. (2005). Effect of short-term cigarette smoke exposure on body weight, appetite and brain neuropeptide Y in mice. Neuropsychopharmacology, 30(4), 713–719. 10.1038/sj.npp.1300597

Cole, R. D., Poole, R. L., Guzman, D. M., Gould, T. J., & Parikh, V. (2015). Contributions of β2 subunit-containing nAChRs to chronic nicotine-induced alterations in cognitive flexibility in mice. Psychopharmacology, 232(7), 1207–1217. 10.1007/s00213-014-3754-4

Copping, N. A., Berg, E. L., Foley, G. M., Schaffler, M. D., Onaga, B. L., Buscher, N., Silverman, J. L., & Yang, M. (2017). Touchscreen learning deficits and normal social approach behavior in the Shank3B model of Phelan–McDermid Syndrome and autism. Neuroscience, 345, 155–165. 10.1016/j.neuroscience.2016.05.016

Cribbs, D. H., Berchtold, N. C., Perreau, V., Coleman, P. D., Rogers, J., Tenner, A. J., & Cotman, C. W. (2012). Extensive innate immune gene activation accompanies brain aging, increasing vulnerability to cognitive decline and neurodegeneration: a microarray study. Journal of Neuroinflammation, 9(179). 10.1186/1742-2094-9-179

Cross, S. J., Linker, K. E., & Leslie, F. M. (2017). Sex-dependent effects of nicotine on the developing brain. Journal of Neuroscience Research, 95(1–2), 422–436. 10.1002/jnr.23878

DeVito, E. E., Herman, A. I., Waters, A. J., Valentine, G. W., & Sofuoglu, M. (2014). Subjective, physiological, and cognitive responses to intravenous nicotine: Effects of sex and menstrual cycle phase. Neuropsychopharmacology, 39(6), 1431–1440. 10.1038/npp.2013.339

Durand, C. M., Betancur, C., Boeckers, T. M., Bockmann, J., Chaste, P., Fauchereau, F., Nygren, G., Rastam, M., Gillberg, I. C., Anckarsäter, H., Sponheim, E., Goubran-Botros, H., Delorme, R., Chabane, N., Mouren-Simeoni, M. C., De Mas, P., Bieth, E., Rogé, B., Héron, D., … Bourgeron, T. (2007). Mutations in the gene encoding the synaptic scaffolding protein SHANK3 are associated with autism spectrum disorders. Nature Genetics, 39(1), 25–27. 10.1038/ng1933

Emilio Merlo Pich, Sonia R. Pagliusi, Michela Tessari, Dominique Talabot-Ayer, Rob Hooft van Huijsduijnen, & Christian Chiamulera. (1997). Common Neural Substrates for the Addictive Properties of Nicotine and Cocaine. Science, 275, 83–86. 10.1126/science.275.5296.83

F. Bovet-Nitti. (1966). Facilitation of Simultaneous Visual Discrimination by Nicotine in the Rat. Psychopharmacologia (BerI.), 10, 59–66.

F. Bovet-Nitti. (1969). Facilitation of Simultaneous Visual Discrimination by Nicotine in Four “Inbred” Strains of Mice. Psychopharmacologia (Berl.), 14, 193–199.

Friedman, N. P., & Robbins, T. W. (2022). The role of prefrontal cortex in cognitive control and executive function. Neuropsychopharmacology, 47(1), 72–89. 10.1038/s41386-021-01132-0

Gotti, C., Zoli, M., & Clementi, F. (2006). Brain nicotinic acetylcholine receptors: native subtypes and their relevance. In Trends in Pharmacological Sciences (Vol. 27, Issue 9, pp. 482–491). 10.1016/j.tips.2006.07.004

Gould, T. J. (2023). Epigenetic and long-term effects of nicotine on biology, behavior, and health. In Pharmacological Research (Vol. 192). Academic Press. 10.1016/j.phrs.2023.106741

Grimm, J. W., Ratliff, C., North, K., Barnes, J., & Collins, S. (2012). Nicotine increases sucrose self-administration and seeking in rats. Addiction Biology, 17(3), 623–633. 10.1111/j.1369-1600.2012.00436.x

Haroon, E., Miller, A. H., & Sanacora, G. (2017). Inflammation, Glutamate, and Glia: A Trio of Trouble in Mood Disorders. Neuropsychopharmacology, 42(1), 193–215. 10.1038/npp.2016.199

Harrod, S. B., Mactutus, C. F., Bennett, K., Hasselrot, U., Wu, G., Welch, M., & Booze, R. M. (2004). Sex differences and repeated intravenous nicotine: Behavioral sensitization and dopamine receptors. Pharmacology Biochemistry and Behavior, 78(3), 581–592. 10.1016/j.pbb.2004.04.026

Hart, E., Hertia, D., Barrett, S. T., & Charntikov, S. (2021). Varenicline rescues nicotine-induced decrease in motivation for sucrose reinforcement. Behavioural Brain Research, 397. 10.1016/j.bbr.2020.112887

He, Z., Xu, Y., Rao, Z., Zhang, Z., Zhou, J., Zhou, T., & Wang, H. (2024). The role of α7-nAChR-mediated PI3K/AKT pathway in lung cancer induced by nicotine. In Science of the Total Environment (Vol. 912). Elsevier B.V. 10.1016/j.scitotenv.2023.169604

Heishman, S. J., Kleykamp, B. A., & Singleton, E. G. (2010). Meta-analysis of the acute effects of nicotine and smoking on human performance. In Psychopharmacology (Vol. 210, Issue 4, pp. 453–469). 10.1007/s00213-010-1848-1

Hogle, J. M., & Curtin, J. J. (2006). Sex differences in negative affective response during nicotine withdrawal. Psychophysiology, 43(4), 344–356. 10.1111/j.1469-8986.2006.00406.x

Hollenhorst, M. I., & Krasteva-Christ, G. (2021). Nicotinic acetylcholine receptors in the respiratory tract. Molecules, 26(20). 10.3390/molecules26206097

Horner, A. E., Heath, C. J., Hvoslef-Eide, M., Kent, B. A., Kim, C. H., Nilsson, S. R. O., Alsiö, J., Oomen, C. A., Holmes, A., Saksida, L. M., & Bussey, T. J. (2013). The touchscreen operant platform for testing learning and memory in rats and mice. Nature Protocols, 8(10), 1961–1984. 10.1038/nprot.2013.122

Horner, A. E., Norris, R. H., McLaren-Jones, R., Alexander, L., Komiyama, N. H., Grant, S. G. N., Nithianantharajah, J., & Kopanitsa, M. V. (2021). Learning and reaction times in mouse touchscreen tests are differentially impacted by mutations in genes encoding postsynaptic interacting proteins SYNGAP1, NLGN3, DLGAP1, DLGAP2 and SHANK2. Genes, Brain and Behavior, 20(1). 10.1111/gbb.12723

Imamura, K., Suzumura, A., Sawada, M., Mabuchi, C., & Marunouchi, T. (1994). Induction of MHC class II antigen expression on murine microglia by interleukin-3. Journal of Neuroimmunology, 55, 119–125.

Jarvik, M. E. (1991). Beneficial effects of nicotine. British Journal of Addiction, 86(5), 571–575. 10.1111/j.1360-0443.1991.tb01810.x

Jias, L. M., & Ellison, G. (1990). Chronic Nicotine Induces a Specific Appetite for Sucrose in Rats. Pharmacology Biochemistry & Behavior, 35(2), 489–491. 10.1016/0091-3057(90)90192-K

Kawakami, E., Nakaoka, S., Ohta, T., & Kitano, H. (2016). Weighted enrichment method for prediction of transcription regulators from transcriptome and global chromatin immunoprecipitation data. Nucleic Acids Research, 44(11), 5010–5021. 10.1093/nar/gkw355

Kim, K., & Picciotto, M. R. (2023). Nicotine addiction: More than just dopamine. In Current Opinion in Neurobiology (Vol. 83). Elsevier Ltd. 10.1016/j.conb.2023.102797

Komiyama, M., Yamakage, H., Satoh-Asahara, N., Ozaki, Y., Morimoto, T., Shimatsu, A., Takahashi, Y., & Hasegawa, K. (2018). Sex differences in nicotine dependency and depressive tendency among smokers. Psychiatry Research, 267, 154–159. 10.1016/j.psychres.2018.06.010

Koylu, E., Demirgören, S., London, D. E., & Pö□ün, S. (1997). Sex difference in up-regulation of nicotinic acetylcholine receptors in rat brain. Life Sciences, 61(12), PL185–PL190. 10.1016/S0024-3205(97)00665-6

Kuleshov, M. V., Jones, M. R., Rouillard, A. D., Fernandez, N. F., Duan, Q., Wang, Z., Koplev, S., Jenkins, S. L., Jagodnik, K. M., Lachmann, A., McDermott, M. G., Monteiro, C. D., Gundersen, G. W., & Maayan, A. (2016). Enrichr: a comprehensive gene set enrichment analysis web server 2016 update. Nucleic Acids Research, 44(1), W90–W97. 10.1093/nar/gkw377

Kumar, M., Keady, J., Aryal, S. P., Hessing, M., Richards, C. I., & Turner, J. R. (2024). The Role of Microglia in Sex- and Region-Specific Blood-Brain Barrier Integrity During Nicotine Withdrawal. Biological Psychiatry Global Open Science, 4(1), 182–193. 10.1016/j.bpsgos.2023.08.019

Lacy, R. T., Hord, L. L., Morgan, A. J., & Harrod, S. B. (2012). Intravenous gestational nicotine exposure results in increased motivation for sucrose reward in adult rat offspring. Drug and Alcohol Dependence, 124(3), 299–306. 10.1016/j.drugalcdep.2012.01.025

Liu, X., Zhan, Z., Li, D., Xu, L., Ma, F., Zhang, P., Yao, H., & Cao, X. (2011). Intracellular MHC class II molecules promote TLR-triggered innate immune responses by maintaining activation of the kinase Btk. Nature Immunology, 12(5), 416–424. 10.1038/ni.2015

Macpherson, T., & Hikida, T. (2018). Nucleus accumbens dopamine D1-receptor-expressing neurons control the acquisition of sign-tracking to conditioned cues in mice. Frontiers in Neuroscience, 12(JUN). 10.3389/fnins.2018.00418

Mcgrath-Morrow, S. A., Gorzkowski, J., Groner, J. A., Rule, A. M., Wilson, K., Tanski, S. E., Collaco, J. M., & Klein, J. D. (2020). The Effects of Nicotine on Development. Pediatrics, 145(3), e20191346. 10.1542/peds.2019-1346

Michael Feyder, Rose-Marie Karlsson, Poonam Mathur, Matthew Lyman, Roland Bock, Reza Momenan, Jeeva Munasinghe, Maria Luisa Scattoni, Jessica Ihne, Marguerite Camp, Carolyn Graybeal, Douglas Strathdee, Alison Begg, Veronica A. Alvarez, Peter Kirsch, Marcella Rietschel, Sven Cichon, Henrik Walter, Andreas Meyer-Lindenberg, … Andrew Holmes. (2010). Association of Mouse Dlg4 (PSD-95) Gene Deletion and Human DLG4 Gene Variation With Phenotypes Relevant to Autism Spectrum Disorders and Williams’ Syndrome. American Journal of Psychiatry, 12(167). 10.1176/appi.ajp.2010.10040484

Milienne-Petiot, M., Higa, K. K., Grim, A., Deben, D., Groenink, L., Twamley, E. W., Geyer, M. A., & Young, J. W. (2018). Nicotine improves probabilistic reward learning in wildtype but not alpha7 nAChR null mutants, yet alpha7 nAChR agonists do not improve probabilistic learning. European Neuropsychopharmacology, 28(11), 1217–1231. 10.1016/j.euroneuro.2018.08.005

Milienne-Petiot, M., Kesby, J. P., Graves, M., van Enkhuizen, J., Semenova, S., Minassian, A., Markou, A., Geyer, M. A., & Young, J. W. (2017). The effects of reduced dopamine transporter function and chronic lithium on motivation, probabilistic learning, and neurochemistry in mice: Modeling bipolar mania. Neuropharmacology, 113, 260–270. 10.1016/j.neuropharm.2016.07.030

Moen, J. K., & Lee, A. M. (2021). Sex Differences in the Nicotinic Acetylcholine Receptor System of Rodents: Impacts on Nicotine and Alcohol Reward Behaviors. In Frontiers in Neuroscience (Vol. 15). Frontiers Media S.A. 10.3389/fnins.2021.745783

Morita, M., Wang, Y., Sasaoka, T., Okada, K., Niwa, M., Sawa, A., & Hikida, T. (2016). Dopamine D2L Receptor Is Required for Visual Discrimination and Reversal Learning. Molecular Neuropsychiatry, 2(3), 124–132. 10.1159/000447970

Myers, C. S., Taylor, R. C., Moolchan, E. T., & Heishman, S. J. (2008). Dose-related enhancement of mood and cognition in smokers administered nicotine nasal spray. Neuropsychopharmacology, 33(3), 588–598. 10.1038/sj.npp.1301425

Neumann, H., Boucraut, J., Hahnel, C., Misgeld, T., & Wekerle, H. (1996). Neuronal control of MHC class II inducibility in rat astrocytes and microglia. European Journal of Neuroscience, 8(12), 2582–2590. 10.1111/j.1460-9568.1996.tb01552.x

Nikiforuk, A., Kos, T., Potasiewicz, A., & Popik, P. (2015). Positive allosteric modulation of alpha 7 nicotinic acetylcholine receptors enhances recognition memory and cognitive flexibility in rats. European Neuropsychopharmacology, 25(8), 1300–1313. 10.1016/j.euroneuro.2015.04.018

Nisell, M., Nomikos, G. G., Chergui, K., Grillner, P., & Svensson, T. H. (1997). Chronic Nicotine Enhances Basal and Nicotine-Induced Fos Immunoreactivity Preferentially in the Medial Prefrontal Cortex of the Rat. In Neuropsychopharmacology (Vol. 17, Issue 3).

Nishioka, T., Attachaipanich, S., Hamaguchi, K., Lazarus, M., de Kerchove d’Exaerde, A., Macpherson, T., & Hikida, T. (2023). Error-related signaling in nucleus accumbens D2 receptor-expressing neurons guides inhibition-based choice behavior in mice. Nature Communications, 14(1). 10.1038/s41467-023-38025-3

Ohmura, Y., Tsutsui-Kimura, I., & Yoshioka, M. (2012). Impulsive behavior and nicotinic acetylcholine receptors. In Journal of Pharmacological Sciences (Vol. 118, Issue 4, pp. 413–422). Japanese Pharmacological Society. 10.1254/jphs.11R06CR

Ortega, L. A., Tracy, B. A., Gould, T. J., & Parikh, V. (2013). Effects of chronic low- and high-dose nicotine on cognitive flexibility in C57BL/6J mice. Behavioural Brain Research, 238(1), 134–145. 10.1016/j.bbr.2012.10.032

Pinto, D., Pagnamenta, A. T., Klei, L., Anney, R., Merico, D., Regan, R., Conroy, J., Magalhaes, T. R., Correia, C., Abrahams, B. S., Almeida, J., Bacchelli, E., Bader, G. D., Bailey, A. J., Baird, G., Battaglia, A., Berney, T., Bolshakova, N., Bölte, S., … Betancur, C. (2010). Functional impact of global rare copy number variation in autism spectrum disorders. Nature, 466(7304), 368–372. 10.1038/nature09146

Pogun, S., Yararbas, G., Nesil, T., & Kanit, L. (2017). Sex differences in nicotine preference. In Journal of Neuroscience Research (Vol. 95, Issues 1–2, pp. 148–162). John Wiley and Sons Inc. 10.1002/jnr.23858

Ponzoni, L., Sala, C., Verpelli, C., Sala, M., & Braida, D. (2019). Different attentional dysfunctions in eEF2K−/−, IL1RAPL1−/− and SHANK3Δ11−/− mice. Genes, Brain and Behavior, 18(5). 10.1111/gbb.12563

Qin, L., Ma, K., Wang, Z. J., Hu, Z., Matas, E., Wei, J., & Yan, Z. (2018). Social deficits in Shank3-deficient mouse models of autism are rescued by histone deacetylase (HDAC) inhibition. Nature Neuroscience, 21(4), 564–575. 10.1038/s41593-018-0110-8

Radyushkin, K., Hammerschmidt, K., Boretius, S., Varoqueaux, F., El-Kordi, A., Ronnenberg, A., Winter, D., Frahm, J., Fischer, J., Brose, N., & Ehrenreich, H. (2009). Neuroligin-3-deficient mice: Model of a monogenic heritable form of autism with an olfactory deficit. Genes, Brain and Behavior, 8(4), 416–425. 10.1111/j.1601-183X.2009.00487.x

Rosecrans, J. A. (1972). Brain area nicotine levels in male and female rats with different levels of spontaneous activity. Neuropharmacology, 11.

Sato, D., Lionel, A. C., Leblond, C. S., Prasad, A., Pinto, D., Walker, S., O’Connor, I., Russell, C., Drmic, I. E., Hamdan, F. F., Michaud, J. L., Endris, V., Roeth, R., Delorme, R., Huguet, G., Leboyer, M., Rastam, M., Gillberg, C., Lathrop, M., … Scherer, S. W. (2012). SHANK1 deletions in males with autism spectrum disorder. American Journal of Human Genetics, 90(5), 879–887. 10.1016/j.ajhg.2012.03.017

Schilström, B., De Villiers, S., Malmerfelt, A., Svensson, T. H., & Nomikos, G. G. (2000). Nicotine-induced fos expression in the nucleus accumbens and the medial prefrontal cortex of the rat: Role of nicotinic and NMDA receptors in the ventral tegmental area. Synapse, 36(4), 314–321. 10.1002/(SICI)1098-2396(20000615)36:4<314::AID-SYN8>3.0.CO;2-U

Schwartz, L. P., Kearns, D. N., & Silberberg, A. (2018). The effect of nicotine pre-exposure on demand for cocaine and sucrose in male rats. Behavioural Pharmacology, 29(4), 316–326. 10.1097/FBP.0000000000000357

Su, W., Sun, J., Shimizu, K., & Kadota, K. (2019). TCC-GUI: A Shiny-based application for differential expression analysis of RNA-Seq count data. BMC Research Notes, 12(1). 10.1186/s13104-019-4179-2

Südhof, T. C. (2008). Neuroligins and neurexins link synaptic function to cognitive disease. In Nature (Vol. 455, Issue 7215, pp. 903–911). Nature Publishing Group. 10.1038/nature07456

Szklarczyk, D., Kirsch, R., Koutrouli, M., Nastou, K., Mehryary, F., Hachilif, R., Gable, A. L., Fang, T., Doncheva, N. T., Pyysalo, S., Bork, P., Jensen, L. J., & Von Mering, C. (2023). The STRING database in 2023: protein-protein association networks and functional enrichment analyses for any sequenced genome of interest. Nucleic Acids Research, 51(1 D), D638–D646. 10.1093/nar/gkac1000

Turner, J. A., Padgett, C., McDonald, S., Ahuja, K. D. K., Francis, H. M., Lim, C. K., & Honan, C. A. (2021). Innate immunity impacts social-cognitive functioning in people with multiple sclerosis and healthy individuals: Implications for IL-1ra and urinary immune markers. Brain, Behavior, and Immunity - Health, 14. 10.1016/j.bbih.2021.100254

Vargas-Medrano, J., Carcoba, L. M., Vidal Martinez, G., Mulla, Z. D., Diaz, V., Ruiz-Velasco, A., Alvarez-Primo, F., Colina, G., Iñiguez, S. D., Thompson, P. M., O’Dell, L. E., & Gadad, B. S. (2023). Sex and diet-dependent gene alterations in human and rat brains with a history of nicotine exposure. Frontiers in Psychiatry, 14. 10.3389/fpsyt.2023.1104563

von Mering, C., Huynen, M., Jaeggi, D., Schmidt, S., Bork, P., & Snel, B. (2003). STRING: A database of predicted functional associations between proteins. In Nucleic Acids Research (Vol. 31, Issue 1, pp. 258–261). 10.1093/nar/gkg034

von Mering, C., Jensen, L. J., Snel, B., Hooper, S. D., Krupp, M., Foglierini, M., Jouffre, N., Huynen, M. A., & Bork, P. (2005). STRING: Known and predicted protein-protein associations, integrated and transferred across organisms. Nucleic Acids Research, 33(DATABASE ISS.). 10.1093/nar/gki005

Warburton, D. M., Rusted, J. M., & Fowler, J. (1992). A comparison of the attentional and consolidation hypotheses for the facilitation of memory by nicotine. Psychopharmacology, 108, 443–447.

West, K. A., Brognard, J., Clark, A. S., Linnoila, I. R., Yang, X., Swain, S. M., Harris, C., Belinsky, S., & Dennis, P. A. (2003). Rapid Akt activation by nicotine and a tobacco carcinogen modulates the phenotype of normal human airway epithelial cells. Journal of Clinical Investigation, 111(1), 81–90. 10.1172/JCI200316147

Wieczorek, M., Abualrous, E. T., Sticht, J., Álvaro-Benito, M., Stolzenberg, S., Noé, F., & Freund, C. (2017). Major histocompatibility complex (MHC) class I and MHC class II proteins: Conformational plasticity in antigen presentation. In Frontiers in Immunology (Vol. 8, Issue MAR). Frontiers Research Foundation. 10.3389/fimmu.2017.00292

Winkler, D., Daher, F., Wüstefeld, L., Hammerschmidt, K., Poggi, G., Seelbach, A., Krueger-Burg, D., Vafadari, B., Ronnenberg, A., Liu, Y., Kaczmarek, L., Schlüter, O. M., Ehrenreich, H., & Dere, E. (2018). Hypersocial behavior and biological redundancy in mice with reduced expression of PSD95 or PSD93. Behavioural Brain Research, 352, 35–45. 10.1016/j.bbr.2017.02.011

Xie, Z., Bailey, A., Kuleshov, M. V., Clarke, D. J. B., Evangelista, J. E., Jenkins, S. L., Lachmann, A., Wojciechowicz, M. L., Kropiwnicki, E., Jagodnik, K. M., Jeon, M., & Ma’ayan, A. (2021). Gene Set Knowledge Discovery with Enrichr. Current Protocols, 1(3). 10.1002/cpz1.90

Zeidler, R., Albermann, K., & Lang, S. (2007). Nicotine and apoptosis. Apoptosis, 12(11), 1927–1943. 10.1007/s10495-007-0102-8

Zoli, M., Pucci, S., Vilella, A., & Gotti, C. (2018). Neuronal and Extraneuronal Nicotinic Acetylcholine Receptors. Current Neuropharmacology, 16(4), 338–349. 10.2174/1570159x15666170912110450

